# Long-term sensitization of rat spinal neurons induced by adolescent psychophysical stress is further enhanced by a mild-nociceptive lumbar input

**DOI:** 10.1101/2021.04.30.442126

**Authors:** Sathish Kumar Singaravelu, Alexander Dawit Goitom, Akseli Surakka, Handan Moerz, Andreas Schilder, Anita C Hansson, Rainer Spanagel, Rolf-Detlef Treede

## Abstract

**Background:** Non-specific low back pain (LBP) is one of the most common chronic pain conditions and adverse childhood experiences (ACEs) are known mediators for chronicity of LBP. Sensitization of dorsal horn neurons (DHNs) is a significant element that contributes to chronic LBP. Repeated restraint stress in adult animals is known to cause manifest DHN sensitization when combined with a short-lasting nociceptive input.

**Objective:** In this study, we investigated whether repeated restraint stress in early adolescence leads to a long-term sensitization of DHNs and if an additional mild-nociceptive input by intramuscular nerve growth factor (NGF) leads to further sensitization.

**Methods:** Adolescent Wistar rats were stressed repeatedly in a narrow plastic restrainer, 1 hour per day for 12 consecutive days. Control animals were handled but not restrained. In adulthood, rats were treated with intramuscular injections of saline or NGF (short-lasting mild-nociceptive input) into the lumbar multifidus muscle (L5). Behavioral tests for pain sensitivity were performed before, after stress and inconjunction with intramuscular injections. Rats were transcardially perfused and immunohistochemistry was performed on lumbar (L2) spinal segments.

**Results:** Adolescent restraint stress significantly lowered the low back pressure pain threshold (PPT) immediately after the stress (*p*<0.0001) and was maintained throughout adulthood (*p*<0.05). Additionally, paw withdrawal threshold (PWT) was significantly lowered by stress (*p*<0.0001) but normalized towards adulthood. An NGF injection in adulthood in previously stressed animals lowered the PPT (Cohen’s *d*=0.87) and increased microglia marker (Iba-1) immunoreactive area in the superficial DHN (*p*<0.01) with a trend in increased feret’s diameter of the immunoreactive cell (*p*=0.05).

**Conclusions:** Our adolescence stress model induced behavioral signs of sensitization and enhanced sensitivity to further sensitization and dorsal horn microglia activation by subsequent mild nociceptive input (NGF injection). These findings help to understand certain aspects of how adolescent stress might predispose to exacerbation of pain with an additional insult.

## 1. Introduction

Adolescence is a critical period in the development and adverse childhood experiences (ACEs) have long-term adverse effects on mental and physical health (Herzog et al., 2018). Furthermore, both animal (Schmidt, 2010) and human (Heim et al., 2010) studies have demonstrated how adolescent stress impacts neuronal circuits and how the immune system development is implicated in enhanced susceptibility of physio- and psychopathological states in adulthood. Specifically, early life stressors such as emotional abuse are associated with enhanced temporal summation of pain and sexual abuse with enhanced touch sensitivity (Tesarz et al., 2016). In humans, stress and ACEs are risk factors for the chronicity of subacute low back pain (LBP) (Mendelek et al., 2013) and the development of chronic widespread pain (Tesarz et al., 2015). However, the mechanistic underpinnings remain elusive and the long-term impact of adolescent stress on adult pain sensitivity remains understudied.

Experimental animal models mimic several human pain conditions (Challa et al., 2015). Stress-related structural or functional changes are likely to impact pain sensitivity (Sandkühler et al., 2009). Previously, we have shown how a single nerve growth factor (NGF) injection into the low back muscle or repeated restraint stress in adulthood can induce a state of latent sensitization in DHNs i.e., the neurons behaved normally but were more easily sensitized by an additional nociceptive input by NGF injection (Hoheisel et al., 2013; 2015). The resulting manifest sensitization was characterized by mechanical hyperalgesia, a significant increase in resting activity, and expansion of receptive fields toward hind limb to deep tissue stimulation (Hoheisel et al., 2013; Singaravelu et al., 2021), all hallmark signs of nociceptive central sensitization.

A wide range of studies indicates that insults across adolescence induce robust and long-lasting effects on pain summation (Burke et al., 2013; Deng et al., 2017; Genty et al., 2018; Le Coz et al., 2017; Vilela et al., 2017). Across this long developmental window, the type and timing of the stressor also have significant effects on the development. For example one of the most common types of early life stressors (pre-weaning) used is the maternal separation that has bidirectional effects on sensitivity to pain in adulthood (Burke et al., 2013; Genty et al., 2018; Vilela et al., 2017). This bidirectionality may be led on by differential microglia activation that has been implicated as a potential mechanism of action for modulating sensitivity to pain and stress (Tsuda et al., 2004; Tynan et al., 2010). Involvement of glial cells in the sensitization of DHNs has also been shown in NGF based animal model of myofascial LBP (Zhang et al., 2017).

Nerve growth factor is produced and released by several cell types (Aloe et al., 2015) and its production is higher in overloaded or injured muscles (Hayashi et al., 2011; Murase et al., 2010; Wu et al., 2009). In humans, intramuscular NGF injections did not elicit immediate pain sensations (Deising et al., 2012) but induced long-lasting hyperalgesia (Svensson et al., 2003, Weinkauf et al., 2015). Furthermore, recent studies have shown that inhibiting NGF alleviated pain in chronic LBP patients (Denk et al., 2017; Kelleher et al., 2016; Leite et al., 2014; Markmann et al., 2020). In mice, the combination of stress and nociceptive intramuscular NGF injections induced manifest hyperalgesia and increased glial cells expression (LaPorta et al., 2020; Lomazzo et al., 2015).

Thus, depending on the stress model and intensity, it seems plausible that prolonged stress or NGF-induced mild nociceptive input in animals may play prominent roles in changing the responsiveness of the neurons in the central nervous system. Nonetheless, to our knowledge, no study have investigated the long-term effect of repeated restraint stress during adolescence on pain related sensitization and their potential neurobiological mechanisms.

In the present study, we hypothesized that repeated restraint stress during adolescence induces a long-term sensitization leading to increased sensitivity to pain stimuli. Additionally, we hypothesized that a subsequent mild-nociceptive input in adulthood further enhances this sensitization to lumbar inputs. For this purpose, we evaluated stimulus-evoked pain-related behaviors first in adolescence and then in adulthood followed by an immunohistological analysis of microglia in L2 spinal cord sections.

## 2. Materials and Methods

### 2.1. Experimental animals and housing conditions

Twenty four male Wistar rats were used in the study. Animals arrived from (ENVIGO, Netharlands) on a postnatal day (PD) 21 and were housed in groups of four in standard macrolon cages (length, width, height: 55×35×20cm). Animals had free access to food and water *ad* libitum and were kept on a normal 12h light/dark cycle. All experimental procedure were approved by the Committe on Animal Care and Use (Regierungspräsidium Karlsruhe, Germany) and were carried out in accordance with German Law on the protection of animals and ethical proposals of the International Association for the Study of Pain. One animal died and their results was excluded from further analyses.

#### 2.1.1. Treatment groups with nerve growth factor and saline injections

Animals were briefly (2 hours) acclimatized to their new home cage in the housing room and 1 hour to the experimental procedure room on arrival. The stress paradigm was induced during the early adolescent phase while the injections of saline/NGF into the multifidus (MF) muscle in the adulthood (Schneider et al., 2013) (see Fig. 1). All animals received two injections at an interval of 5 days. Animals were randomly assigned to four treatment groups and each cage had two control and two stress rats. The experimenter was blinded to the treatments the animals received. All the experiments were conducted in the inactivate phase of the animals. The perfusion and tissue collection were performed within 5 h from the start of the inactive cycle.

**Figure 1:**
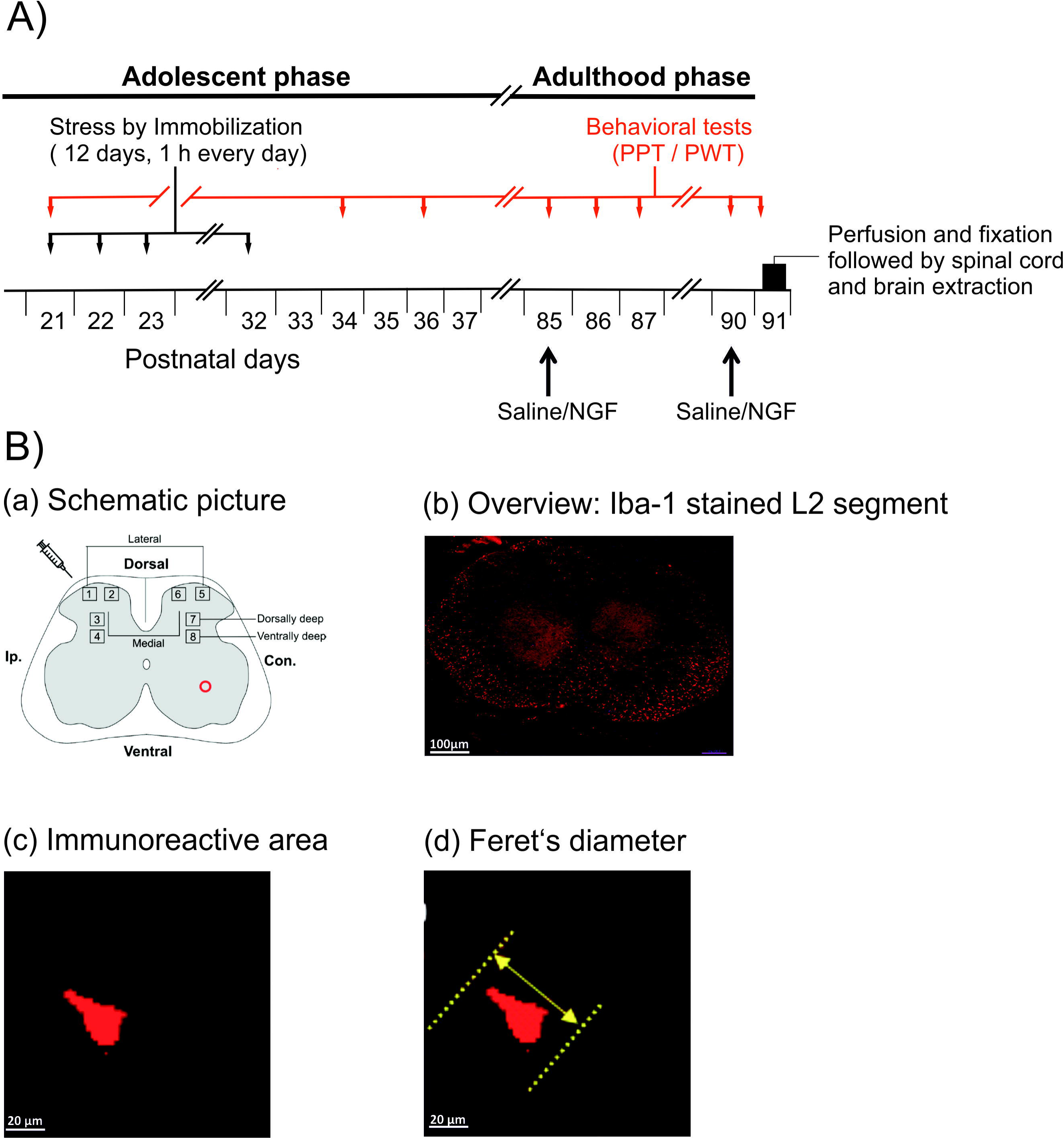
Experimental methods. **A)** Animals of the stress group were repeatedly stressed in a narrow plastic restrainer for 12 consecutive days for 1 h every day on postnatal days (PD21-32; adolescent phase). The pressure pain threshold (PPT) of the multifidus muscle at vertebral level L5 and paw withdrawal threshold (PWT) of the hind paw was measured at different time points (red line). All animals received injections of saline/NGF on PD85 and PD90 (adulthood phase) respective to the group they belonged to. On PD91 (black bar) all animals were perfused and fixed. The spinal cord and brain tissues were extracted and stored. **B.a)** a schematic illustration of the region of interests on a spinal lumbar 2 (L2) section (refer section: 2.9.2). **B.b)** example overview image of an L2 section stained for Iba-1. **B.c)** example image for an Iba-1 immunoreactive area calculation (refer section: 2.9.2). **B.d)** example image for an Iba-1 stained cell feret’s diameter calculation (refer section: 2.9.2).

#### 2.1.1.1. Control + saline + nerve growth factor (CSN)

The control animals (n=6) were handled on 12 consecutive days (transported to the laboratory and manipulated by hand) similar to the stress animals without repeated restraint stress. These animals received saline as their 1^st^ injection followed by NGF at an interval of 5 days in adulthood.

#### 2.1.1.2. Stress + saline + saline (RSS)

Repeated restraint stress (n=6) was induced on 12 consecutive days for 1 hour every day in a narrow plastic tube (refer section 2.2). These animals received 2 saline injections 5 days apart.

#### 2.1.1.3. Control + nerve growth factor + nerve growth factor (CNN)

The positive controls (n=6) and were handled similar to stress animals without repeated restraint stress. These animals received 2 NGF injections at a 5 day interval in adulthood.

#### 2.1.1.4. Stress + saline + nerve growth factor (RSN)

Repeated restraint stress (n=5) was induced on 12 consecutive days for 1 hour every day in a narrow plastic tube (refer section 2.2). These animals received saline as their 1^st^ injection followed by NGF at an interval of 5 days in adulthood.

### 2.2. Repeated restraint stress (R)

Restraint stress was induced as previously described (Hoheisel et al., 2015) by placing animals in a narrow restrainer (inner length 15 cm; inner height 4 cm, Fig.1) for 1 h daily on 12 consecutive days. Body weight was measured on 12 consecutive days before the stress paradigm and on days before the behavioral experiment (refer section 2.4). We observed no differences in body weight between stressed and control animals during or after the 12 days stress paradigm or in adulthood. Signs of distress such as vocalization, struggling during restraint (escape movements), urination, and/or defecation were observed during the stress paradigm.

### 2.3. Injection of nerve growth factor and saline

As a second intervention animals received injections of saline and NGF or both at two different time points with an interval of 5 days (see Fig. 1). Injections of 50 µl NGF at a concentration of 0.8 µM (NGF, human recombinant, Calbiochem, Merck, Germany) dissolved in phosphate buffer saline (PBS: pH of the NGF solution 7.2-7.3) were administered into the left multifidus muscle 3 mm lateral to the spinous process at vertebral level L5 (Hoheisel et al., 2013; Zhang et al., 2017). The NGF concentration used is known to induce hyperalgesia when intramuscularly injected in animals or humans (Deising et al., 2012; Hoheisel et al., 2013; Hoheisel et al., 2007; Svensson et al., 2003). The second injections were administered to the same site. Injections of 50 µl saline (vehicle) at a concentration of 0.9% served as control. No visible signs of muscle inflammation were observed after the injections of neither NGF nor saline (Hoheisel et al., 2013).

### 2.4 Behavioral tests

Behavioral tests were performed to assess for mechanical allodynia/hyperalgesia. All behavioral measurements were performed by the same experimenter and the experimental setups were cleaned with 70% ethanol between animals. See timeline of the behavioral experiments (redline in Fig.1).

### 2.5 Pressure pain threshold

To test for mechanical hyperalgesia the pressure pain threshold (PPT) of the low back, was determined with an electronic Von Frey anaesthesiometer (Life Science Instruments, Woodland Hills, CA, USA). A blunt tip with an area of 3.46 mm^2^ was pressed with increasing intensity to the MF muscle through the intact skin at the vertebral level L5. With the blunt tip, mainly nociceptors in deep tissues but not in the skin were excited (Takahashi et al., 2005). The PPT was defined as the minimum pressure intensity that is required to elicit a pain-related reaction (withdrawal and escape movements, vocalization).

### 2.6 Paw withdrawal threshold

The paw withdrawal sensitivity to mechanical stimuli was measured with an electronic von Frey esthesiometer equipped with a rigid cylindrical tip (diameter 0.8mm^2^) (Electronic von Frey esthesiometer, IITC Inc. Life Science, USA). After the PPT measurement, the animals were placed into a Plexiglas box (length, width, height: 20 × 10 × 14 cm) with a metal grid as a base for a further 30 min of habituation. Using the electronic esthesiometer, pressure was applied to the plantar region of both the hind paws (ipsilateral to the injection site followed by contralateral) until the rat withdrew it. The paw withdrawal threshold (PWT) is calculated as the mean of five independent measurements. Before the first measurement on PD 36, the animals were habituated to the metal grid table on two consecutive days (PD 34 and 35) for 45 min. The von Frey test measures nociception by provoking the nocifensive hind-paw flexion reflex with mechanical stimulation. This reflex corresponds to the human lower limb flexor reflex and does not equal the complexity of feeling pain. However, in humans, the reflex intensity correlates well with pain perception (Sandrini et al., 2005). This test is valuable to monitor changes at the peripheral and spinal levels of the somatosensory system.

### 2.7 Perfusion

One day after the 2nd injection of NGF or saline, the animals were killed with an overdose of thiopental sodium i.p. (Trapanal®, Inresa GmbH, Germany) and transcardially perfused with 4% paraformaldehyde (PFA) in 0.1 M PBS. Transcardial perfusion started with 0.1 M PBS for 1 – 2 min. The clearing of the liver was used as an indicator of good perfusion. When the liver was clear, 4% PFA was perfused for 20-30 min, followed by 20-30 min PBS perfusion.

### 2.8 Tissue processing

After the perfusion, a laminectomy was performed and the spinal segment L2 was removed and stored in 10% sucrose solution in 0.1 M PBS at 4° C for 1 day. One day before the freezing, spinal segments were transferred to a 30% sucrose solution in 0.1 M PBS at 4° C. The tissues were rapidly frozen on dry ice. The 30% sucrose solution served as a cryoprotectant, dehydrated the tissue, and prevented the formation of ice crystal artifacts in the frozen tissue section. A 20 µm thick cross-section of the spinal segments was made on a cryostat (Cryostat NX70, Theromo Fischer Scinetific Inc., USA) and mounted on glass slides. For immunofluorescence staining, three cross-sections of the spinal segment L2 were selected from each animal at random and each group had five animals.

### 2.9 Immunofluorescence labeling, image processing, and quantification

#### 2.9.1 Ionized Calcium-Binding (Iba-1) labeling for microglia

Structural changes of microglial cells were visualized by Iba-1 immunohistochemistry. Iba-1 is a protein that is specifically expressed in microglia and is upregulated during the activation of microglia (Hinwood et al., 2012; Romero-Sandoval et al., 2008a). The sections were first incubated in 10 % Roti®-block (Carl Roth, Germany) at room temperature for 1 h. Then they were incubated in rabbit anti-Iba-1 polyclonal antibody (1:1000; Abcam, United Kingdom) at room temperature for 16 h. Afterward, sections were incubated in Cy3™ - conjugated goat-anti-rabbit IgG antibody (1:500; Dianova, Germany) at room temperature in dark for 4 h. The Cy3™ - conjugated secondary antibody was detected by a DPSS laser (Leica, Germany) at 561 nm.

#### 2.9.2 Image processing and quantification

Digitized images were obtained with Leica Scanning Microscope (Leica TCS SP8 AOBS confocal laser-scanning microscope, Leica Microsystems, Wetzlar, Germany) using 10x and 40x oil immersion objective lens with a computer-based imaging software LAS-AF (Leica, Germany). Immunofluorescence was acquired using a scanning sequential mode to avoid crosstalk among simultaneously scanned channels with two laser lines: Argon – laser (488 nm) and DPSS – laser (561 nm) (Leica Microsystems, Germany). Three-dimensional images were acquired over 20-μm z-axis with a 1.0 - μm step size. All the images were prepared in a maximum intensity Z-stack projection.

For quantitative measurements, 8 regions of interest (ROIs) were selected from the dorsal horn of every section. They were ipsilateral (Ip.) and contralateral (Con.) to the injection site. At each side, 4 ROIs of 255 μm × 255 μm were defined, 2 of which were located in the superficial dorsal horn (medial and lateral; laminae I - II); 2 at various depths in the deep dorsal horn (dorsally and ventrally deep; laminae IV – V; see Fig. 1.B.a). All images were measured with identical imaging parameters. The images were evaluated with an image analysis software Image J (NIH, USA).

An example of an overview image for Iba-1 stained L2 spinal segment is shown in Fig. 1.B.b. For the immunoreactive area, the relative total area was calculated as the percentage of the total immunoreactive area within the ROI divided by the area of the ROI and is expected to increase with hypertrophism of cells (Chacur et al., 2009) and unspecific background was adjusted for each animal (see Fig. 1.B.c). Feret’s diameter is the longest distance between two parallel lines perpendicular to that distance (longest distance) and drawn at the boundaries of the immunoreactive area (see Fig. 1B.d).

### 2.10 Data analysis

Before calculations, the data of PPT and PWT were transformed into decadic logarithms to achieve secondary normal distribution (Bartlett MS., 1947), since previous data obtained in larger cohorts provided solid evidence for log-normal distribution of PPT and other psychophysical data (Rolke et al., 2005). The log PPT and PWT values were then normalized to baseline (log value – log baseline value) for further analysis (equivalent to calculating percentage changes in PPT and PWT).

The PPT and PWT data after stress are shown as individual values with their respective median (Fig. 2.A and 2.B). For the immunoreactive area and feret’s diameter (Fig. 4.A and 4.B), data are shown as mean ± SEM. The *U*-test of Mann and Whitney was used to compare, between and within the groups. A probability level of less than 5% (*p*<0.05, two-tailed, statistical software: GraphPad Prism) was regarded as significant. The PPT and PWT data after saline/NGF injections are shown as individual paired values (Fig. 3.A and 3.C) and effect sizes were determined using Cohen *d* (difference in means divided by pooled SD). An effect size >0.2 was considered as ‘small,’ >0.5 as ‘medium,’ and >0.8 as ‘large’ (Cohen J., 1969).

**Figure 2:**
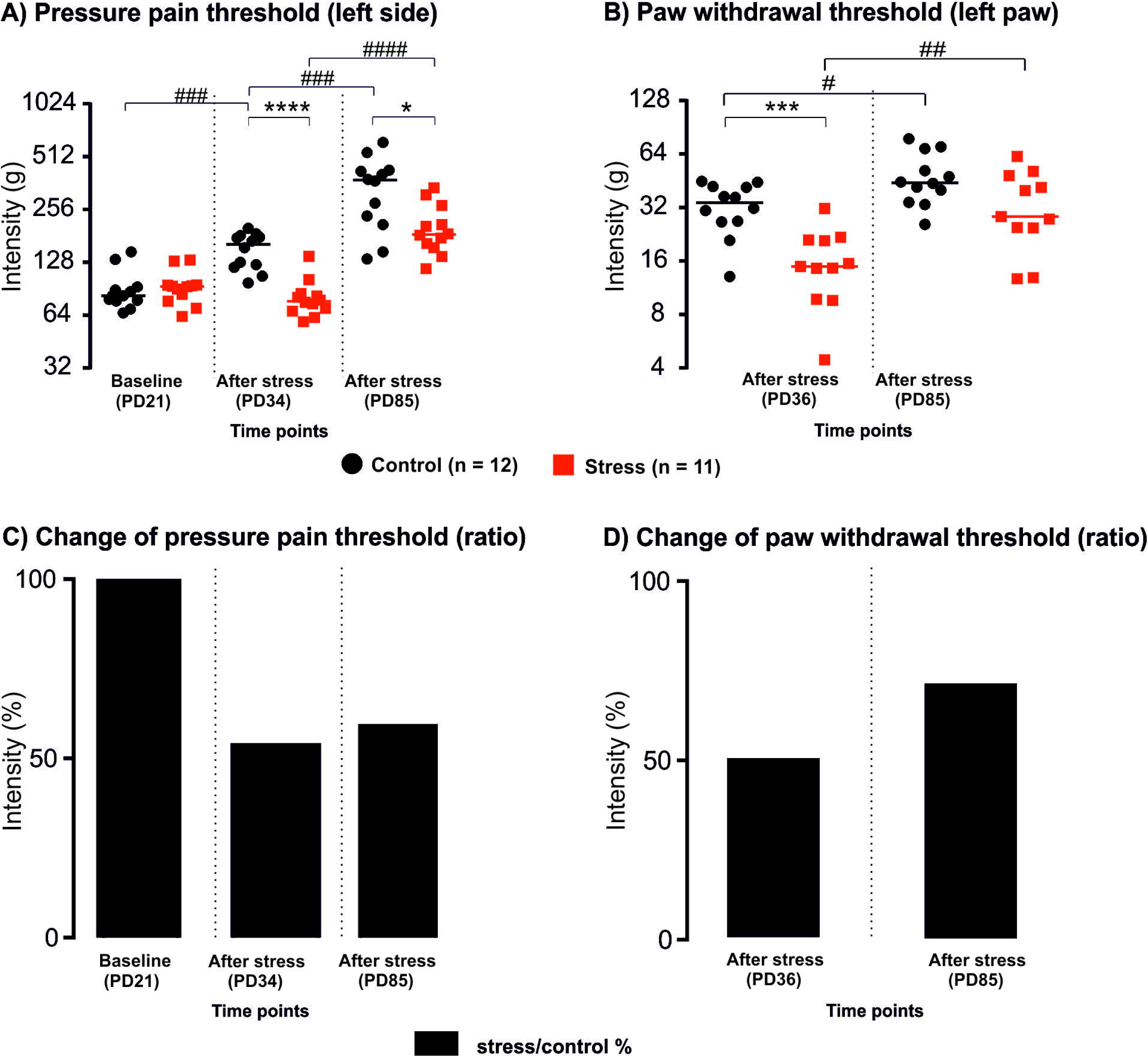
Pressure pain threshold and paw withdrawal threshold before and after stress. **A)** Data expressed in log scale, force (in ‘g’ on the left y-axis) required to elicit a pain-related reaction (withdrawal behavior, escape movements, vocalization) using a blunt probe with an area of 3.46 sq.mm when applied to the multifidus muscle of the low back. Horizontal lines indicate the median for each group. **B)** Data expressed in log scale, force (in ‘g’ on the left y-axis) required to elicit a pain-related reaction (paw licking, paw withdrawal) using a rigid cylindrical tip with an area of 0.8 sq.mm when applied to the plantar surface of the hind paw. Horizontal lines indicate the median for each group. **C)** Intensity (in ‘%’), change of pressure pain threshold is shown in ratio (stress/control). **D)** Intensity (in ‘%’), change of paw withdrawal threshold is shown in ratio (stress/control). Black horizontal lines in (A) and (B) indicate the median for each group. *P*-values: *U*-test of Mann and Whitney; *p*<0.05 is represented with (*/#), *p*<0.0063 (**/##), *p*<0.0001 (***/###), *p*<0.0001 (****/####).

**Figure 3:**
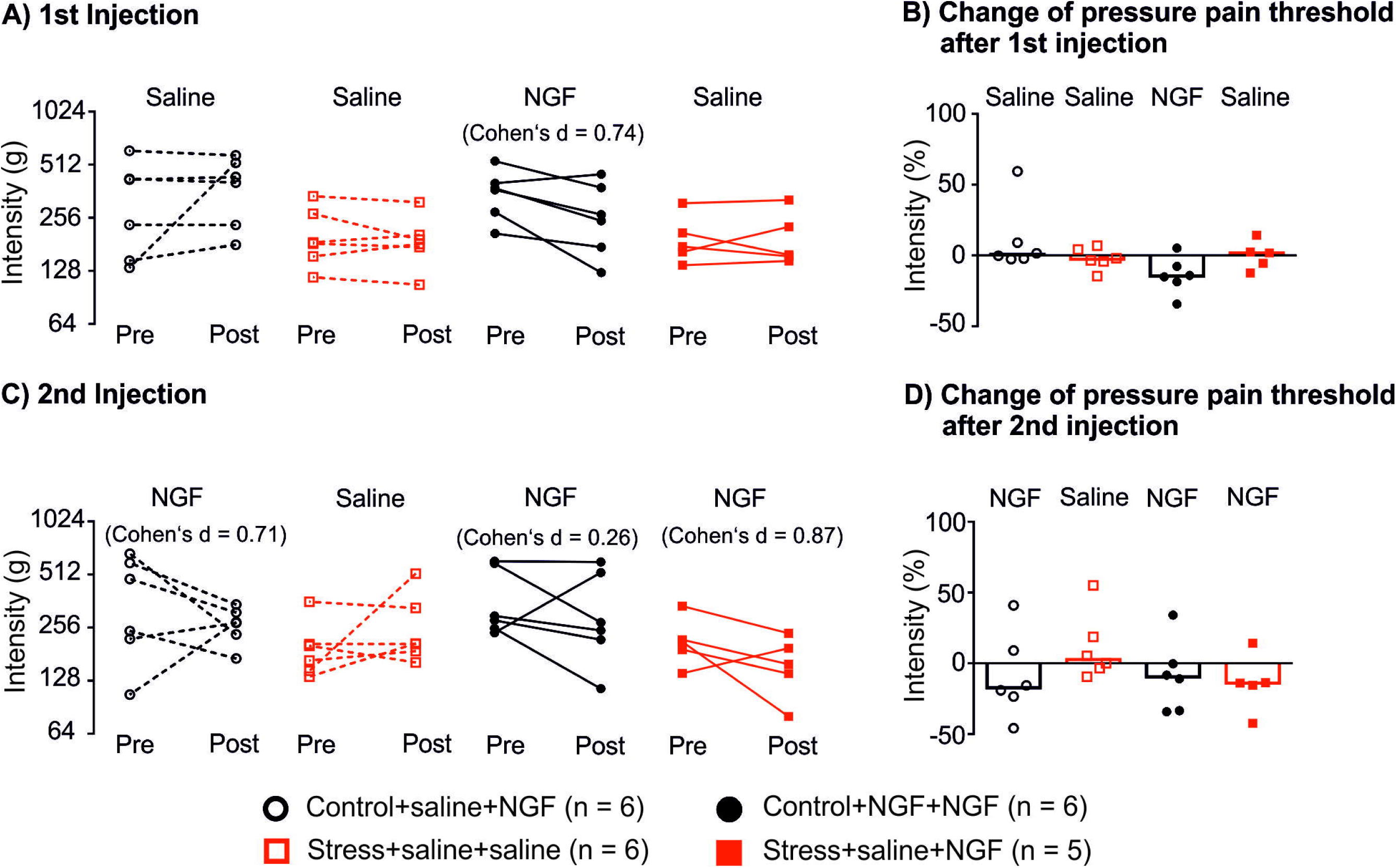
Pressure pain threshold and paw withdrawal threshold before and after saline/NGF injection. **A)** First injection individual animal data expressed in log scale, force (in ‘g’ on the left y-axis) required to elicit a pain-related reaction (withdrawal behavior, escape movements, vocalization) using a blunt probe with an area of 3.46 sq.mm when applied to the multifidus muscle of the low back. **B)** Intensity of the pressure pain threshold expressed in ratio log(post-injection - pre-injection) after the 1^st^ injection. Bars indicate the median for each group. **C)** Second injection individual animal data expressed in log scale, force (in ‘g’ on the left y-axis) required to elicit a pain-related reaction (withdrawal behavior, escape movements, vocalization) using a blunt probe with an area of 3.46 sq.mm when applied to the multifidus muscle of the low back. **D)** Intensity of the pressure pain threshold expressed in ratio log(post-injection - pre-injection) after the second injection. Bars indicate the median for each group.

## 3. Results

### 3.1 Pressure pain threshold after stress

The baseline measurement (PD21) for the PPT of the multifidus muscle revealed no significant difference between the control and stress groups. Two days after the stress paradigm (PD34), the PPT significantly dropped (*p*<0.0001) in the stress group, suggesting that the stress paradigm induced hyperalgesia and this sensitization was still significant (*p*<0.05) in adulthood (PD85). The PPT increase within groups is led by age and is indicated using ‘#’ (Fig. 2. A). In Fig. 2.C., see the change in PPT shown in a ratio (stress/control) before and after stress.

#### 3.1.1 Paw withdrawal threshold after stress

Four days after the stress paradigm (PD36), the PWT significantly lowered (*p*<0.0001) in the repeated restraint stress group, suggesting that the stress paradigm induced hyperalgesia but this sensitization did not continue into the adulthood phase (PD85). These data suggest that the stress paradigm induced a short-term central sensitization (Fig. 2. B). The PWT increase within groups refers to anincrease in age and is indicated using ‘#’ (Fig. 2. B). In Fig. 2.D., one could see the change in PWT shown in a ratio (stress/control) after stress.

### 3.2 Pressure pain threshold of the low back after saline/NGF injections

Intramuscular injections of saline/NGF were administered to each animal as a second intervention to induce sensitization and the animals were divided into four groups (see Sec: 2.1.1). In Fig.3.A. a paired plot of the PPT between pre and post 1^st^ injection of either saline or NGF is shown for individual animals. The group that received NGF injection showed a decrease in PPT with medium effect size (CNN: Cohen’s d = 0.74) while the groups that received saline showed a small or no effect. In Fig. 3.B., the % change of PPT is shown after the 1^st^ injection of saline or NGF in log (post-injection/pre-injection). The group that received NGF shows a change in the PPT in the negative direction implying a drop in the PPT after NGF administration.

In Fig.3.C. a paired plot of the PPT between pre and post 2^nd^ injection of either saline or NGF is shown for individual animals. The groups that received NGF injections showed a decrease in PPT with high effect size (CSN: Cohen’s d = 0.71; RSN: Cohen’s d = 0.87) while the group that received repeated NGF injection showed only a small effect size (CNN: Cohen’s d =0.26). In Fig. 3.D., the % change of PPT is shown after the 2^nd^ injection of saline or NGF in log (post-injection/pre-injection). All the groups that received NGF showed a change in the PPT in the negative direction implying a drop in the PPT after NGF administration.

#### 3.2.1 Paw withdrawal threshold after saline/NGF injections

No effects were observed in the PWT after the injections of saline/NGF. This implies that NGF induced mechanical hyperalgesia is local to the site of injection.

### 3.3 Repeated restraint stress or NGF leads to the increased immunoreactive area and feret’s diameter of Iba-1 stained cells in the spinal dorsal horn neurons

To study the contribution of glial cell markers in the stress+NGF or NGF alone induced sensitization of spinal DHNs, we performed immunohistochemistry on the spinal cord sections.

In the superficial DHN (lamina I – II), we found a significant increase (*p*<0.05) in the immunoreactive area in the groups that received at least one NGF injection compared to the group that received only saline injections (Fig 4.A). In the NGF group that was preceded by stress, the increase in immune reactive area was significant (RSS vs. RSN; *p*<0.01) compared to saline group preceded by stress (Fig.4.A). This demonstrates that a single NGF injection is sufficient to increase the Iba-1 stained cell immunoreactive area. But when preceded by stress, the observed highly significant effect might explain the difference between latent vs. manifest sensitization. In deep DHN, the latent vs. manifest significance exists only between RSS vs. CNN (*p*<0.05). This illustrates how repeated NGF injections can alter the morphology of glial cells in the deep DHN whereas single NGF only altered the superficial DHN microglia when preceded by stress, this alteration becomes highly significant. No significant changes were observed on the contralateral site of injection.

**Figure 4:**
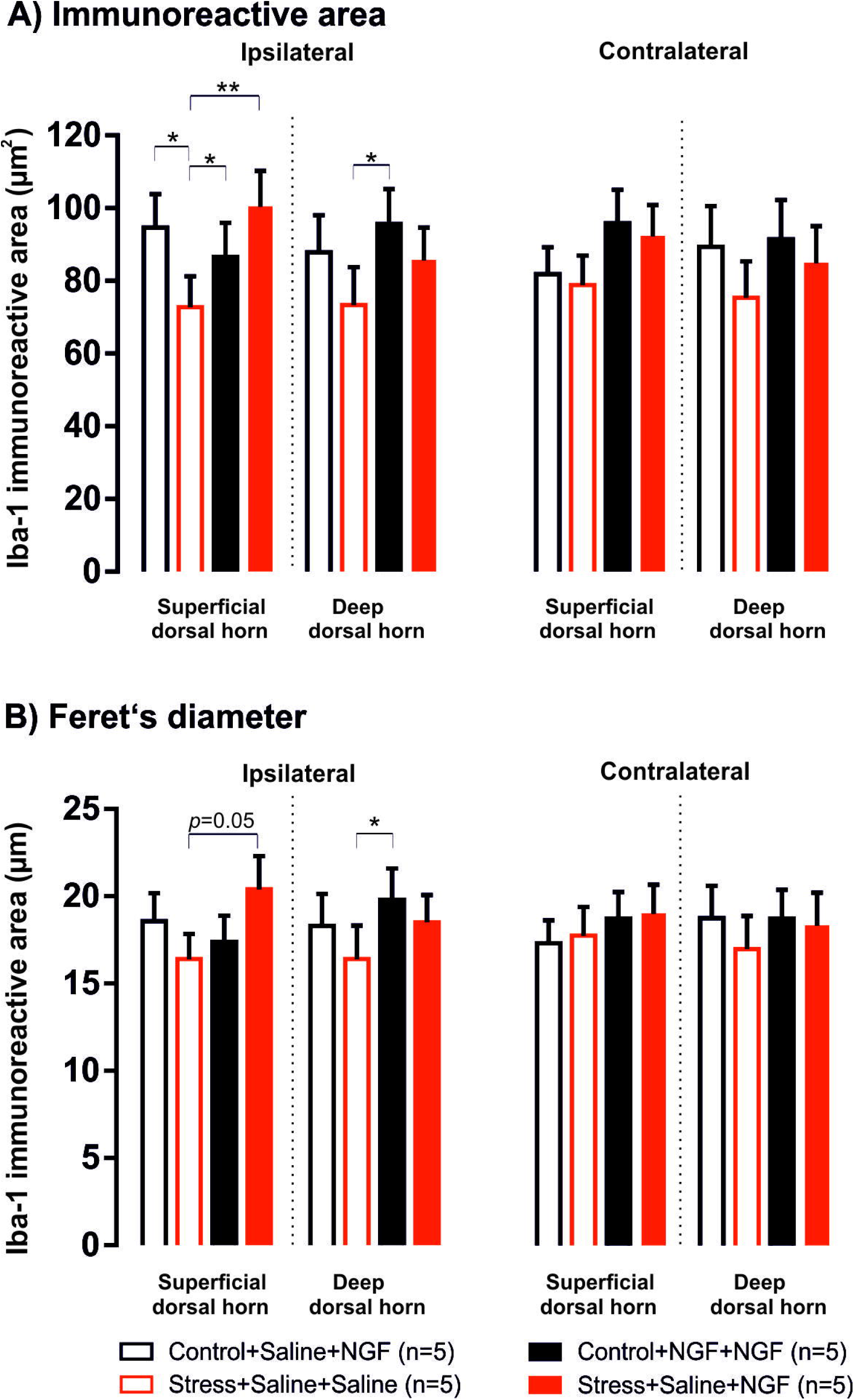
Immunohistochemistry data of stained microglial cells. In each animal, three tissue sections were imaged and analyzed and each group had five animals. Injection of nerve growth factor (NGF) in groups that previously experienced stress or NGF, leads to increased expression of **A)** Immunoreacive area of microglial cells to the ipsilateral side of the injection and **B)** increased Feret’s diameter also to the ipsilateral side of the injection. The inset in the right top corner illustrates the 8 ROI’s (see Sec: 2.9.2), the red circle in the inset indicates the manually made pinhole to identify the contralateral site. *P*-values: *U*-test of Mann and Whitney. *p*<0.05 is represented with (*), *p*<0.01 (**).

In Fig. 4.B., there is a trend in increased feret’s diameter between RSS vs. RSN (*p*=0.05) in the superficial DHN and a significant increase in the deep DHN between the RSS vs. CNN (*p*=0.05). This data further indicates that stress+NGF or repeated NGF increases feret’s diameter of the immunoreactive cells which could contribute to manifest sensitization of the DHNs. No significant changes were observed in the contralateral side of injection.

## Discussion

The present study indicates that psychophysical restraint stress across early and mid adolescence increases sensitivity to painful stimulation of the low back well into adulthood in male Wistar rats. Furthermore, we demonstrated that adolescent restraint stress in conjunction with NGF injection in adulthood induced pain related changes and alterations to microglia morphology.

### 4.1 Adolescent pain sensitivity after stress

We observed an increased sensitivity to PPT in the lower back of repeated restraint stress (RRS) animals shortly after the cessation of the stress paradigm (PD34) (Fig 2.). Furthermore, increased sensitivity was also observed in the PWT compared to controls. These data are indicative of RRS inducing acute sensitization to both deep and cutaneous nociceptive inputs. In line with our findings are reports of other types of stressor such as; social instability stress inducing hypersensitization to pain (Le Coz et al., 2017). Interestingly, they found that social stress led to a modulation of cold reponsiveness, but not mechanical stimulation. Together our results suggest that different stressors modulate pain manifestation differently.

### 4.2 Adult pain sensitivity after adolescent stress

Surprisingly, no studies have followed up on potential long-term consequecnes of adolescent stress on pain. To better understand the long-term effects of adolescent RRS on pressure pain modulation, we reassessed animals in the PPT and PWT again in adulthood (PD85). RRS rats demonstrated increased sensitivity to pressure pain (PPT) over low back muscles when compared to controls. Intresteingly, the difference in pain threshold between groups remained over time, implying that RRS animals show persistent alterations to mechanical stimuli into adulthood. These results expand on previous finding on the effects of early life stress (i.e maternal separation). Where, maternal separation appears to increase mechanical sensivity, although results vary. One study found an increase in mechanical pain sensitivity in adulthood across sexes (Vilela et al., 2017), while others have found mechanical allodynia only in females (Burke et al., 2013). This variability in findings is most likely due to the differences in experimental design as well and strain differences that have been reported.

We observed no significant difference between groups in the PWT in adulthood. Together these data suggest that the observed sensitization to cutaneous input in the PWT after the termination of adolescent stress was acute. However, the sensitization to deep inputs (PPT) seems to be maintained into adulthood in RRS animals, on top of the age related increase in pressure pain threshold as the animals grew (Fig 2.). Since stress did not lead to injuries in low back or paws, this sensitization is likely mediated centrally by enhanced responsiveness of dorsal horn neurons; central sensitization of signal processing in other CNS regions involved in the behavioral responses may also play a role. Considering that LBP is the most common form on pain experienced in human, our findings present a novel model to investigate the long-term effects of adolescent stress on pain modulation. Wistar rats appear to be suitable for such studies, as they are known to be more sensitive to stressors than other strains (López-Rubalcava et al., 2000)

### 4.3 Latent vs Manifest sensitization long-after stress and nociceptive inputs

Results from our current experiment indicate that adolescent RRS alone was strong enough to induce long-lasting sensitization to nociceptive input from the lower back as observed in the PPT test in adulthood (Fig 2.). It is therefore likely that the sensitization may well be manifest and a second stimuli would further exacerbate pain related outcomes. Similar to our previous findings, we observed that an initial NGF injection increased sensivity to pressure pain in the PPT on control animals and this hypersensitivity returned to baseline on day two in Wistar rats, as observed in our previous studies in Sprague-dawley rats (Hoheisel et al., 2013; Zhang et la., 2017). In all cases the second NGF injection led to increased sensitivity in the PPT in both stressed and control animals, with stressed animals maintaining a significantly lower threshold when compared to controls. In other words, single NGF induces short term sensitization but when preceded by stress, it turns into manifest sensitization, as observed in the PPT. An exacerbated effect of repeated NGF was not observed in this study on Wistar rats, unlike previous studies in Sprague-Dawley rats. But like in Spargue-Dawley rats, the effect of manifest sensitization can be seen in the morphological changes of microglia both in the CNN and RSN groups compared to RSS (Fig. 4).

### 4.4 Persistent effects of repeated restraint stress and NGF on microglia morphology

Our findings demonstrate that adolescent stress in conjunction with an NGF injection in adulthood or repeated NGF injections increased microglial immunoreactive area and the feret’s diameter.

All groups that received NGF as the second injection showed a increase in the microglia immunoreactivity area on the inpsilateral side compared to saline injections; there were no contralateral changes. This may implay increased migration of microglia to undertake their phagocygotic state, whether the state of change is neuroprotective or neurotoxic phenotype remains elusive. Evidence suggests that the microglia in the NGF groups are becoming phagocytic as this phenotype is typically associated with increases in size (immunoreactive area) and roundness (feret’s diameter) (Tan et al., 2020). Prior studies have shown region specific difference on a microscale in the brain (Tynan et al., 2010). Although we did not analyze specific regions, this raises an interesting question, suggesting there may be region specific difference in the spinal cord. Our finding provide partial evidence on the interaction of stress and NGF on microglia function and further assessment of additional microglia parameters are needed to characterize the microglia phenotype. However, our results nonetheless improve our understanding on the interaction of adolescent stress and later pain sensitivity

### 4.5 Limitations

Although our findings provide new important evidence on the long-term impact of adolescent stress and the effects of a later “double-hit” on pain sensitivity and microglia morphology, there are still a few aspects that need to be considered. First, strain differences in pain sensitivity and stress sensitivity have been reported; Wistar rats were chosen because of their known stress sensitivity and the observed effects of stress were substantial; future studies should use milder adolescent stressors in order to be sensitive to additive effects of subsequent mild nociceptive input by NGF injection (Singaravelu et al. 2021; Zhang et al., 2003). Second, we lack a pure negative control (control + saline + saline) as a comparison group, this limits the interpretation of the results and should be considered in future studies. Third, due to the setup of the experimental design we were unable to obtain a baseline measurement for the PWT (refer section 2.6). Finally, we assessed only male rats in our study and cannot therefore draw generalizable conclusion on the effects of adolescent RRS on female pain sensivity.

### 4.6 Conclusions

The current data suggest that repeated restraint stress in adolescence induces a long-term central sensitization and primes for enhanced manifest sensitization by a subsequent NGF injection. This enhanced manifest sensitization was indicated by a drop in PPT of the low back muscles, increased immunoreactive area and feret’s diameter of spinal microglia. Particularly, the effect of NGF is more pronounced when it is repeated or preceded by stress and is facilitated by activated glial cells. The stress-induced sensitization of DHNs following NGF injection appears to be of particular importance for the translational from the experimental model presented herein to the clinical condition of musculoskeletal pain, as the increased sensitivity of muscle nociceptors to inflammatory mediators may contribute to the pathophysiology of clinical low back pain.

## Acknowledgments

The authors wish to thank U. Hortscht for her excellent help and assistance in all experiments. The authors acknowledge the support of the Core Facility Live Cell Imaging Mannheim at Mannheim Center for Translational Neuroscience (MCTN) and the Heidelberg Pain Consortium SFB 1158.

